# Synthesis, Structure and Anticancer Activity of a Dinuclear Organoplatinum(IV) Complex Stabilized by Adenine

**DOI:** 10.1101/2024.02.14.580361

**Authors:** Alisha M. O’Brien, William A. Howard, Kraig A. Wheeler

## Abstract

The dinuclear organoplatinum(IV) compound {Pt(CH_3_)_3_}_2_(μ-I)_2_(μ-adenine) (abbreviated **Pt**_**2**_**ad**), obtained by treating cubic [Pt(CH_3_)_3_(μ_3_-I)]_4_ with two equivalents of adenine, was isolated and structurally characterized by single crystal X-ray diffraction. The National Cancer Institute Developmental Therapeutics Program’s *in vitro* sulforhodamine B assays showed **Pt**_**2**_**ad** to be particularly cytotoxic against central nervous system cancer cell line SF-539, and human renal carcinoma cell line RXF-393. Furthermore, **Pt**_**2**_**ad** displayed some degree of cytotoxicity against non-small cell lung cancer (NCI-H522), colon cancer (HCC-2998, HCT-116, HT29, and SW-620), melanoma (LOX-IMVI, MALME-3M, M14, MDA-MB-435, SK-MEL-28, and UACC-62), ovarian cancer (OVCAR-5), renal carcinoma (A498), breast cancer (BT-549 and MDA-MB-468), and triple-negative breast cancer (MDA-MB-231).

## Introduction

FDA-approved platinum-based chemotherapy agents such as cisplatin, carboplatin, and oxaliplatin are the clinical first-line standard of care for epithelial ovarian cancer^1^ and non-small cell lung cancer^2,3^ and often play significant roles in treating other types of cancer as well.^4^ The anticancer effectiveness arises from the interaction between the platinum compound and the cancerous DNA^5,6^ – although alternative mechanisms involving platinum-protein interactions likely contribute to the cytotoxicity as well.^7^ In order to facilitate the interaction with DNA, some experimental platinum-based anticancer drugs have been fitted with flat, aromatic ancillary ligands capable of engaging in π – π interactions with the nucleobases.^8,9,10,11^ These π – π interactions drive the intercalation of the platinum compound into the major or minor grooves of the DNA and bring the platinum center within bonding proximity of the nucleobases.

In light of the significance of such π – π interactions, platinum compounds having nucleobase ligands would seem to be ideal anticancer agents, since the nucleobase ligand would naturally intercalate between base pair layers in the DNA. Indeed, various platinum(II) complexes of nucleotides,^12^ nucleosides,^13,14,15^ and nucleobase derivatives^16^ have been reported as highly lethal to different kinds of cancers. Several platinum(II) compounds having adenosine^17,18^ or an adenine derivative^19,20,21,22,23,24,25,26,27,28,29,30^ as an ancillary ligand have also been reported to be highly cytotoxic toward various cancer cell lines. Although several platinum(IV) complexes with adenine derivatives as ligands have been reported,^31^ we are unaware of any that have been tested as anticancer drugs.

Our research group recently reported the cytotoxicity of octahedral Pt^IV^(CH_3_)_2_I_2_{2,2’-bipyridine} against the human breast cancer cell line ZR-75-1 by the *in vitro* MTT assay method.^32^ This compound is easily prepared by treating [Pt(CH_3_)_2_I_2_]_x_ with 2,2’-bipyridine. Indeed, the Lewis acids [Pt(CH_3_)_2_X_2_]_x_ (X = Br, I) and cubic [Pt(CH_3_)_3_(μ_3_-I)] 4^33^ can be treated with diverse Lewis base ligands to produce a variety of organoplatinum(IV) derivatives, that could potentially display a broad spectrum of very different anticancer properties. In this work, we report the synthesis, structural characterization, and *in vitro* anticancer activity of {Pt(CH_3_)_3_}_2_(μ-I)_2_(μ-adenine), a compound prepared by treating [Pt(CH_3_)_3_(μ_3_-I)]_4_ with adenine.

## Experimental

### General Considerations

All ^1^H, ^13^C, and ^195^Pt NMR spectra were obtained at room temperature on a Bruker Ascend 600 MHz FTNMR spectrometer running Topspin 3.6 at the frequencies 600.16 MHz, 150.91 MHz, and 129.015 MHz, respectively. ^1^H and ^13^C chemical shifts are reported in parts per million relative to SiMe_4_ (δ = 0) and were referenced internally with respect to the protio solvent impurity (δ = 3.31 ppm for HD_2_COD) or the ^13^C resonances (δ = 49.15 ppm for CD_3_OD), respectively. ^195^Pt NMR spectra were referenced externally to a solution of K_2_PtCl_4_ in D_2_O (δ = – 1620 ppm).^34^ Infrared spectra were recorded as KBr pellets on a Nicolet Magna-IR 560 spectrometer. Elemental analyses were carried out by Atlantic Microlab, Inc. (Norcross, GA). Unless otherwise noted, all reactions and manipulations were carried out in the presence of air. All reagents and solvents were obtained from commercial suppliers and were used without further purification. [Pt(CH_3_)_3_(μ_3_-I)]_4_ was prepared by a procedure modified from that by Clark and Manzer.^33^

### Synthesis of [Pt(CH_3_)_3_(μ_3_-I)]_4_

A solution consisting of [1,5-cyclooctadiene]Pt(CH_3_)_2_ (0.653 g, 1.96 mmol) in CH_3_I (6 mL) in a glass bomb was stirred magnetically at 70°C for 4.5 hours. The pale yellow homogeneous solution was cooled to room temperature and pale yellow crystals precipitated. Yield = 0.370 g (51%). The crystals were spectroscopically identical to [Pt(CH_3_)_3_(μ_3_-I)]_4_ and were used without any further purification.

### Synthesis of {*fac*-Pt(CH_3_)_3_}_2_(μ-I)_2_(μ-adenine)

A mixture of [Pt(CH_3_)_3_(μ_3_-I)]_4_ (0.311 g, 0.212 mmol) and adenine (0.058 g, 0.429 mmol) in CHCl_3_/CH_3_OH (1:1 by volume, 10 mL) was stirred at 75°C for 4 hours. The volatile components were removed *in vacuo*, and the residue was washed with CHCl_3_ (2 × 5 mL), extracted into CH_3_OH (10 mL), and crystallized from the CH_3_OH solution at room temperature as a white powder (yield = 0.135 g). A second batch of the product was crystallized from the CHCl_3_ washes (yield = 0.156 g). Yield = 0.291 g (79 %). Anal. Calc. for C_11_H_23_I_2_N_5_Pt_2_: C, 15.20%; H, 2.67%; N, 8.06%. Found: C, 15.30%; H, 2.86%; N, 7.94%. ^1^H NMR (CD_3_OD, δ ppm): 1.396 ppm (s, 6 H, ^2^J_Pt-H_ = 75 Hz), 1.399 ppm (s, 6 H, ^2^J_Pt-H_ = 75 Hz), 1.68 ppm (s, 3 H, ^2^J_Pt-H_ = 74 Hz), 1.76 ppm (s, 3 H, ^2^J_Pt-H_ = 74 Hz), 8.22 ppm (s, 1 H, ^3^J_Pt-H_ = 13 Hz), 8.51 ppm (s, 1 H, ^3^J_Pt-H_ = 10 Hz). ^13^C{^1^H} NMR (CD_3_OD, δ ppm): –8.04 ppm (^1^J_Pt-C_ = 659 Hz), –7.16 ppm (^1^J_Pt-C_ = 661 Hz), 10.19 ppm (^1^J_Pt-C_ = 708 Hz), 11.68 ppm (^1^J_Pt-C_ = 718 Hz), 112.4 ppm, 144.0 ppm, 154.3 ppm, 155.8 ppm, 157.0 ppm. ^195^Pt{^1^H} NMR (CD_3_OD, δ ppm): – 2863 ppm, – 2929 ppm. IR (KBr): 3442 (vs), 2963 (m), 2911 (m), 2898 (s), 2853 (m),2816 (m), 1653 (s), 1560 (m), 1541 (m), 1509 (w), 1507 (w), 1472 (m), 1457 (m), 1397 (m), 1322 (w), 1266 (w), 1224 (m), 1109 (m), 1082 (m), 1028 (w), 911 (w), 789 (m), 732 (m), 668 (m), 611 (s), 560 (s), 467 (m).

### X-Ray Diffraction Studies

Pale yellow plates of {*fac*-Pt(CH_3_)_3_}_2_(μ-I)_2_(μ-adenine) were crystallized by the slow addition of CHCl_3_ (g) to a C_2_H_5_OH solution at room temperature. X-ray intensity data were collected using a Bruker D8 Venture diffractometer equipped with a graphite monochromator and a Mo Kα micro-focus INCOATEC Iμs 3.0 sealed tube at 0.71073 Å. Data sets were corrected for Lorentz and polarization effects as well as absorption. The criterion for observed reflections is *I > 2σ(I)*. Lattice parameters were determined from least squares analysis and reflection data. Empirical absorption corrections were applied using SADABS.^35^ The structure was solved by direct methods and refined by full-matrix least squares analysis on *F*^*2*^ using X-Seed^36^ equipped with SHELXT.^37^ All non-hydrogen atoms were refined anisotropically by full-matrix least squares on *F*^*2*^ using the SHELXL^38^ program. Hydrogen atoms were included in idealized geometric positions with *U*_*iso*_ *= 1.2 U*_*eq*_ of the atom to which they are attached (*U*_*iso*_ *= 1.5 U*_*eq*_ for methyl groups). Those hydrogen atoms attached to nitrogen or oxygen were located in difference maps and assigned 1.2 x *U*_*eq*_. The frames were integrated with the Bruker SAINT software package using a narrow-frame algorithm. The structure was solved and refined using the Bruker SHELXTL Software Package, and the cell data and refinement parameters are summarized in Table 1. Crystallographic data have been deposited with the Cambridge Crystallographic Data Centre: the deposition number is CCDC 2298069. These data can be obtained free of charge at https://www.ccdc.cam.ac.uk/ or from the Cambridge Crystallographic Data Centre, 12, Union Road, Cambridge CB2 1EZ, UK; fax: +44 (0)1223-336408.

**Table 1.** Crystal and intensity collection data for {fac-Pt(CH_3_)_3_}_2_(μ-I)_2_(μ-adenine) ·CHCl_3_ · CH_3_CH_2_OH.

|  |  |
| --- | --- |
| **formula** | $\text{C}_{14}\text{H}_{30}\text{Cl}_3\text{I}_2\text{N}_5\text{OPt}_2$ |
| **molecular weight, g mol<sup>-1</sup>** | 1034.76 |
| **temperature, K** | 100.00 |
| **wavelength, Å** | 0.71073 |
| **lattice** | triclinic |
| **space group** | $P_{-1}$ |
| **cell constants** |  |
| **a, Å** | 10.1669(9) |
| **b, Å** | 10.5929(9) |
| **c, Å** | 13.0266(11) |
| **α, deg.** | 92.574(3) |
| **β, deg.** | 109.885(3) |
| **γ, deg.** | 103.182(4) |
| **volume, Å<sup>3</sup>** | 1272.97(19) |
| **Z** | 2 |
| **ρ (calc.) g cm<sup>-3</sup>** | 2.700 |
| **absorption coefficient, mm<sup>-1</sup>** | 13.732 |
| **F(000)** | 940 |
| **crystal size, mm<sup>3</sup>** | 0.337 x 0.189 x 0.066 |
| **θ range** | 1.992 to 33.778° |
| **index ranges** | -15 ≤ h ≤ 15<br>-16 ≤ k ≤ 16<br>-20 ≤ l ≤ 20 |
| **reflections collected** | 83,153 |
| **independent reflections** | 10,210 [$R_{\text{int}} = 0.0328$] |
| **coverage, independent reflections** | 100% |
| **Absorption correction** | Multi-scan |
| **max. & min. transmission** | 0.7467 and 0.3892 |
| **refinement method** | Full-matrix least-squares on $F^2$ |
| **data/restraints/parameters** | 10,210/1/263 |
| **goodness-of-fit on $F^2$** | 1.085 |
| **Final R indices [$I > 2\sigma(I)$]** | $R_1 = 0.0223$ $wR_2 = 0.0460$ |
| **R indices (all data)** | $R_1 = 0.0282$ $wR_2 = 0.0475$ |
| **largest difference peak and hole** | 1.413 & -1.998 e. Å<sup>-3</sup> |

### *In Vitro* Sulforhodamine B Assays

*In vitro* assays were carried out by staff members in the National Cancer Institute’s Developmental Therapeutics Program.^39,40,41^ Sixty human tumor cell lines were grown in RPMI 1640 medium containing 5% fetal bovine serum and 2 mM L-glutamine. Cells were inoculated into 96 well microtiter plates in 100 μL at plating densities ranging from 5,000 to 40,000 cells per well, depending on the doubling time of individual cell lines. After cell inoculation, the microtiter plates were incubated at 37°C, 5% CO_2_, 95% air and 100% relative humidity for 24 hours prior to addition of experimental drugs. After 24 hours, 2 plates of each cell line were fixed *in situ* with trichloroacetic acid, to represent a measurement of the cell population for each cell line at the time of drug addition (T_z_). **Pt**_**2**_**ad** was solubilized in N,N-dimethylformamide at 400-fold the desired final maximum test concentration and stored frozen prior to use. At the time of drug addition, an aliquot of frozen concentrate was thawed and diluted to twice the desired final maximum test concentration with a complete medium containing 50 μg/mL gentamicin. An additional four, 10-fold or ½ log serial dilutions were made to provide a total of 5 drug concentrations plus control. 100 μL aliquots of these different drug dilutions were added to the appropriate microtiter wells already containing 100 μL of medium, resulting in the required final drug concentrations. Following drug addition, the plates were incubated for an additional 48 hours at 37°C, 5% CO_2_, 95% air and 100% relative humidity. For adherent cells, the assay was terminated by the addition of cold trichloroacetic acid. Cells were fixed *in situ* by the gentle addition of 50 μL of cold 50% (w/v) trichloroacetic acid (final concentration, 10% trichloroacetic acid) and incubated for 60 minutes at 4°C. The supernatant was discarded, and the plates were washed 5 times with tap water and air-dried. Sulforhodamine B solution (100 μL) at 0.4% (w/v) in 1% acetic acid was added to each well, and plates were incubated for 10 minutes at room temperature. After staining, unbound dye was removed by washing 5 times with 1% acetic acid and the plates were air-dried. Bound stain was subsequently solubilized with 10 mM trizma base, and the absorbance was read on an automated plate reader at wavelength 515 nm. For suspension cells, the methodology was the same, except that the assay was terminated by fixing settled cells at the bottom of the wells by gently adding 50 μL of 80% trichloroacetic acid (final concentration, 16% trichloroacetic acid). Using the 7 absorbance measurements [time zero (T_z_), control growth (C), and test growth in the presence of drug at the 5 concentration levels (T_i_)], the percentage growth was calculated at each of the drug concentration levels. Calculations were carried out as follows:

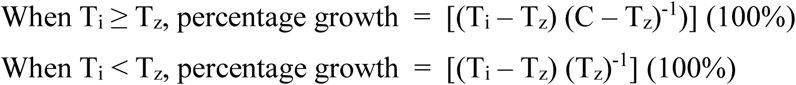

Three dose response parameters were calculated for each experimental agent. Growth inhibition of 50% (GI_50_) was calculated from …..

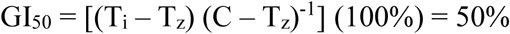

which was the drug concentration resulting in a 50% reduction in the net protein increase (as measured by sulforhodamine B staining) in control cells during the drug incubation. The drug concentration resulting in total growth inhibition (TGI) was calculated from T_i_ = T_z_. The LC_50_ (concentration of drug resulting in a 50% reduction in the measured protein at the end of the drug treatment as compared to that at the beginning) indicating a net loss of cells following treatment was calculated from ….

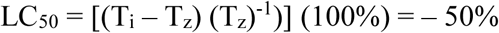

Values were calculated for each of these three parameters if the level of activity was reached; however, if the effect were not reached or were exceeded, the value for that parameter was expressed as greater or less than the maximum of minimum concentration tested.

## Results and Discussion

The new compound {*fac*-Pt(CH_3_)_3_}_2_(μ-I)_2_(μ-adenine) (abbreviated **Pt**_**2**_**ad**) was prepared by treating [Pt(CH_3_)_3_(μ_3_-I)]_4_ with two equivalents of adenine as shown in Figure 1 and isolated as an air-stable crystalline solid. The two platinum atoms in **Pt**_**2**_**ad** are chemically inequivalent, and so the ^195^Pt{^1^H} NMR spectrum features two peaks at δ – 2863 and – 2929 ppm. Four sets of methyl peaks appear with their ^195^Pt satellites in the ^1^H NMR spectrum, at δ 1.396 ppm (6H, ^2^J = 75 Hz), 1.399 ppm (6H, ^2^J = 75 Hz), 1.68 ppm (3H, ^2^J = 74 Hz), and 1.76 ppm (3H, ^2^J = 74 Hz). Likewise, the ^13^C{^1^H} NMR spectrum reveals 4 sets of methyl groups with their ^195^Pt satellites at δ –8.04 ppm (^1^J = 659 Hz), –7.16 ppm (^1^J = 661 Hz), 10.19 ppm (^1^J = 708 Hz), and 11.68 ppm (^1^J = 718 Hz).

**Figure 1.**
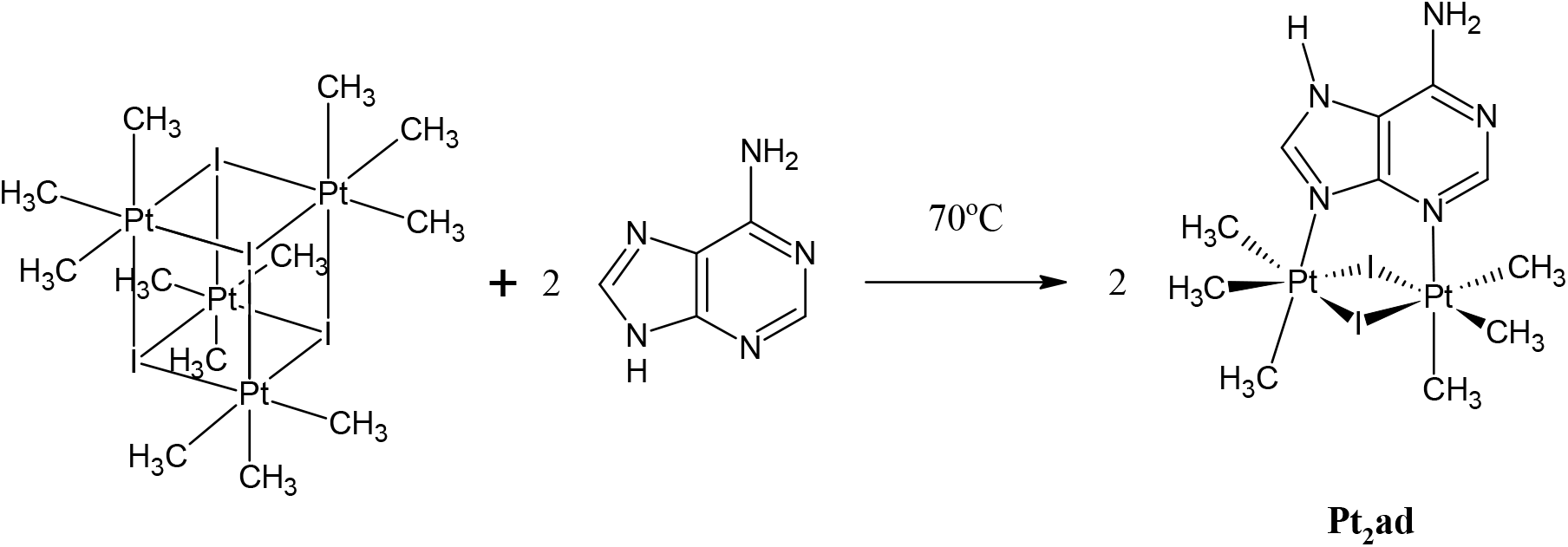
Synthesis of {*fac*-Pt(CH_3_)_3_}_2_(μ-I)_2_(μ-adenine) (**Pt**_**2**_**ad**).

**Pt**_**2**_**ad** was structurally characterized by single crystal X-ray diffraction. Figure 2 shows a thermal ellipsoid plot (50% probability) with the non-hydrogen atoms labeled. Table 1 shows the crystal and intensity collection data, while Table 2 shows select metrical data.

**Table 2.**
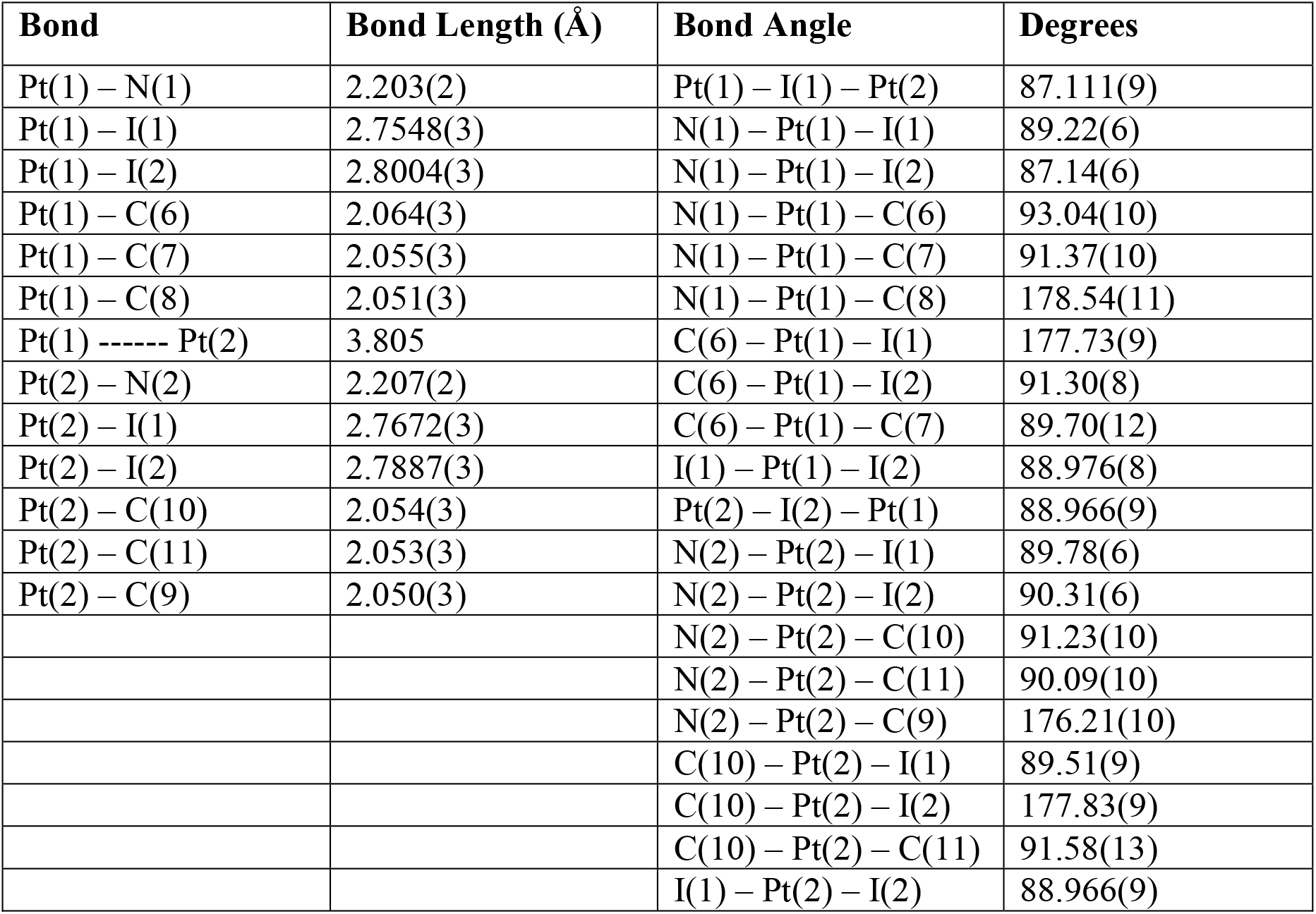
Select Metrical Data.

| Bond | Bond Length (Å) | Bond Angle | Degrees |
| --- | --- | --- | --- |
| Pt(1) – N(1) | 2.203(2) | Pt(1) – I(1) – Pt(2) | 87.111(9) |
| Pt(1) – I(1) | 2.7548(3) | N(1) – Pt(1) – I(1) | 89.22(6) |
| Pt(1) – I(2) | 2.8004(3) | N(1) – Pt(1) – I(2) | 87.14(6) |
| Pt(1) – C(6) | 2.064(3) | N(1) – Pt(1) – C(6) | 93.04(10) |
| Pt(1) – C(7) | 2.055(3) | N(1) – Pt(1) – C(7) | 91.37(10) |
| Pt(1) – C(8) | 2.051(3) | N(1) – Pt(1) – C(8) | 178.54(11) |
| Pt(1) ----- Pt(2) | 3.805 | C(6) – Pt(1) – I(1) | 177.73(9) |
| Pt(2) – N(2) | 2.207(2) | C(6) – Pt(1) – I(2) | 91.30(8) |
| Pt(2) – I(1) | 2.7672(3) | C(6) – Pt(1) – C(7) | 89.70(12) |
| Pt(2) – I(2) | 2.7887(3) | I(1) – Pt(1) – I(2) | 88.976(8) |
| Pt(2) – C(10) | 2.054(3) | Pt(2) – I(2) – Pt(1) | 88.966(9) |
| Pt(2) – C(11) | 2.053(3) | N(2) – Pt(2) – I(1) | 89.78(6) |
| Pt(2) – C(9) | 2.050(3) | N(2) – Pt(2) – I(2) | 90.31(6) |
|  |  | N(2) – Pt(2) – C(10) | 91.23(10) |
|  |  | N(2) – Pt(2) – C(11) | 90.09(10) |
|  |  | N(2) – Pt(2) – C(9) | 176.21(10) |
|  |  | C(10) – Pt(2) – I(1) | 89.51(9) |
|  |  | C(10) – Pt(2) – I(2) | 177.83(9) |
|  |  | C(10) – Pt(2) – C(11) | 91.58(13) |
|  |  | I(1) – Pt(2) – I(2) | 88.966(9) |

**Figure 2.**
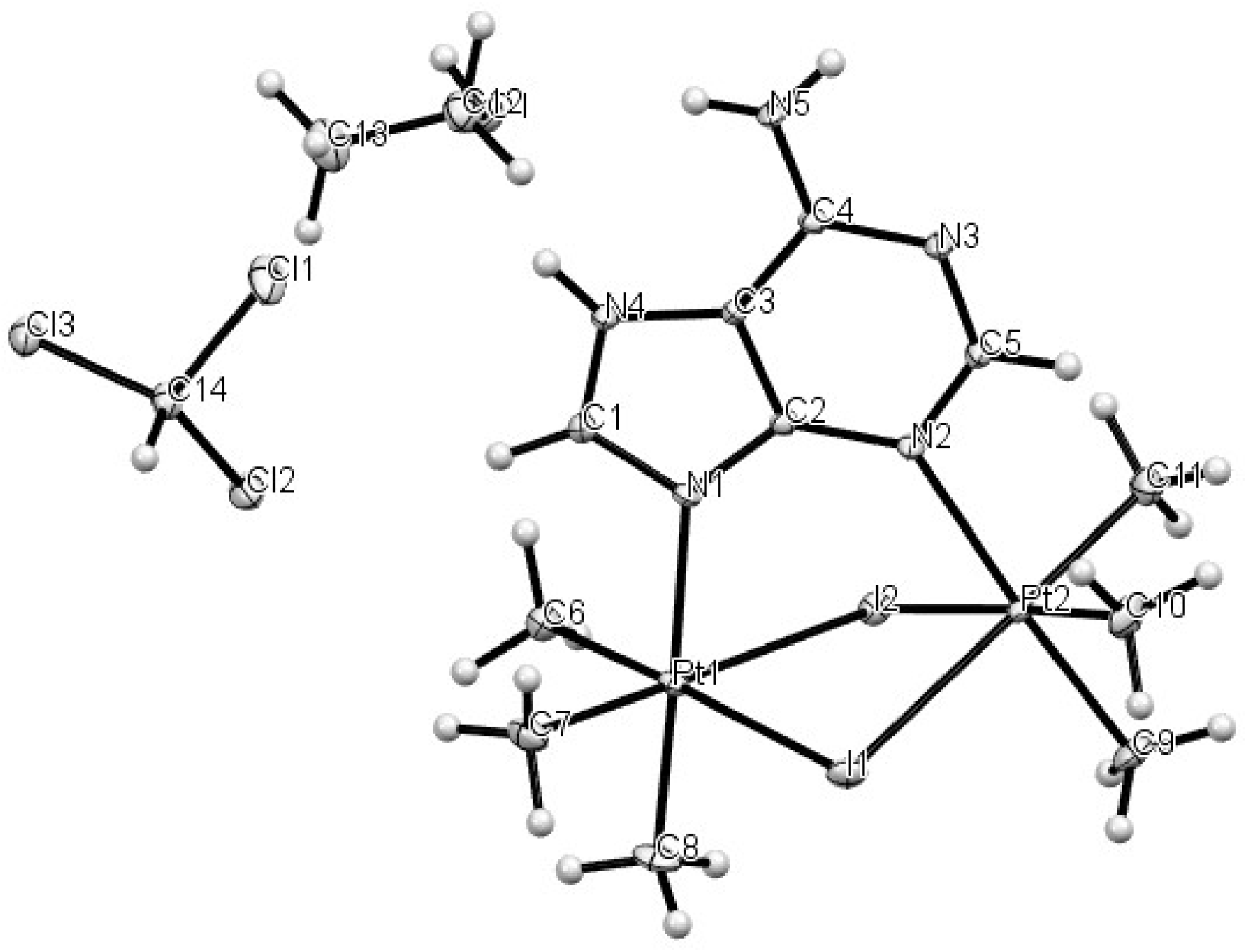
Thermal Ellipsoid Plot (50%) from the X-ray Structure of {*fac*-Pt(CH_3_)_3_}_2_(μ-I)_2_(μ-adenine) · CHCl_3_ · CH_3_CH_2_OH.

Adenine is known to coordinate to both platinum(II) and platinum(IV) through a variety of diverse coordination modes.^31^ The particular coordination mode in **Pt**_**2**_**ad** involves bridging two Pt(IV) centers by a neutral tautomer of adenine, as shown in Figure 2. A similar coordination mode was proposed for {Pt^IV^Cl_3_(H_2_O)}_2_(μ-Cl){μ-adenine(−1)} where adenine(−1) is a deprotonated adenine anion, but this compound was not structurally characterized by X-ray crystallography.^42^

The coordination geometry for each Pt(IV) center in **Pt**_**2**_**ad** is octahedral. The plane defined by I(1) – Pt(1) – I(2) intersects the plane defined by I(1) – Pt(2) – I(2) at an angle of 32.49°. The 3-center, 4-electron Pt – I bond lengths in **Pt**_**2**_**ad** are intermediate between the terminal 2-center, 2-electron Pt – I bonds in Pt(CH_3_)_2_I_2_{2,2’-bipyridine} (2.6355(5) and 2.6569(5) Å),^32^ and the 4-center, 6-electron bonds in cubic [Pt(CH_3_)_3_(μ_3_-I)]_4_ · ½ CH_3_I (2.8105(8) to 2.8366(7) Å).^43^ The Pt – N bond distances in **Pt**_**2**_**ad** closely resemble those in trinuclear [{Pt^IV^(CH_3_)_3_(μ-9-methyladenine(−1))}_3_] · O=C(CH_3_)_2_ (2.18(1) – 2.25(1) Å) and in trinuclear [{Pt^IV^(CH_3_)_3_(μ-9-methyladenine(−1))}_3_] · 1.5 Et_2_O ·2 H_2_O (2.180(7) – 2.226(7) Å).^44^ The Pt(1) ---- Pt(2) distance in **Pt**_**2**_**ad** (3.805 Å) is within the sum of the van der Waals radii of two platinum atoms,^45^ suggesting an intramolecular electronic interaction between the two platinum atoms.

The National Cancer Institute’s Developmental Therapeutics Program (NCI/DTP) tested the anticancer activity of **Pt**_**2**_**ad** against 60 cancer cell lines by the *in vitro* sulforhodamine B assay. More specifically, there were 6 leukemia, 9 non-small cell lung cancer, 7 colon cancer, 6 central nervous system cancer, 9 melanoma, 7 ovarian cancer, 8 renal cancer, 2 prostate cancer, and 6 breast cancer cell lines. **Pt**_**2**_**ad** exhibited cytotoxicity toward some cancer cell lines, but not all. The results involving only those cell lines effected by **Pt**_**2**_**ad** are summarized in Table 3. For comparison, the results involving cisplatin against the same cell lines are also included in Table 3, as values in parentheses.

**Table 3.** In Vitro Sulforhodamine B Assay Results for **Pt**_**2**_**ad**. Values in Parentheses are Results for Cisplatin.

| Type of Cancer | Cell Line | GI<sub>50</sub>, μM | TGI, μM | LC<sub>50</sub>, μM |
| --- | --- | --- | --- | --- |
| Non-Small Cell Lung | NCI-H522 | 1.51 | 3.70 | 18.6 |
| Colon Cancer | HCC-2998 | 13.1 (10.7) | 31.7 (25.5) | 76.8 (60.8) |
|  | HCT-116 | 3.43 (9.24) | 15.5 (> 100) | 52.5 (> 100) |
|  | HT29 | 5.99 (7.81) | 21.2 (> 100) | 63.6 (> 100) |
|  | SW-620 | 3.37 (2.72) | 11.4 (> 100) | 39.2 (> 100) |
| Central Nervous System | SF-539 | 1.47 (0.600) | 2.76 (7.67) | 5.20 (> 100) |
| Melanoma | LOX IMVI | 3.85 (0.961) | 18.4 (7.93) | 58.2 (> 100) |
|  | MALME-3M | 1.95 (1.85) | 8.19 (4.58) | 39.2 (> 100) |
|  | M14 | 2.08 (1.71) | 14.8 (7.68) | 63.4 (35.1) |
|  | MDA-MB-435 | 6.19 (2.77) | 18.4 (14.4) | 46.2 (42.7) |
|  | SK-MEL-28 | 5.10 (2.99) | 17.4 (12.8) | 42.4 (99.5) |
|  | UACC-62 | 3.18 (1.34) | 15.7 (9.39) | 42.0 (37.2) |
| Ovarian Cancer | OVCAR-5 | 2.35 (4.38) | 11.9 (46.4) | 35.2 (> 100) |
| Renal Cancer | A498 | 19.7 (12.7) | 37.8 (26.4) | 72.6 (54.9) |
|  | RXF-393 | ----- | 2.63 (4.58) | 5.99 (24.2) |
| Breast Cancer | MDA-MB-231 | 1.62 (28.1) | 8.79 (> 100) | 60.2 (> 100) |
|  | BT-549 | 3.56 (3.36) | 16.5 (44.9) | 61.2 (> 100) |
|  | MDA-MB-468 | 1.49 (0.233) | 3.85 (0.744) | 19.2 (4.50) |

These *in vitro* cell viability assays clearly underscore the importance of using multiple cell lines of the same type of cancer. For instance, **Pt**_**2**_**ad** was particularly lethal toward central nervous system (CNS) cancer cell line SF-539 with LC_50_ = 5.20 μM, but **Pt**_**2**_**ad** was completely inactive toward five other CNS cancer cell lines (SF-268, SF-295, SNB-19, SNB-75, and U251). For comparison, cisplatin was inactive toward all 6 CNS cancer cell lines. The SF-539 cell line was derived from glioblastoma multiforme cells located in the right, temporoparietal region of the brain of a 34 year-old white female in 1987.^46^ Glioblastoma multiforme is the deadliest of all brain cancers. Combined with radiotherapy, temozolomide^47^ is a first-line standard of care chemotherapy drug for treating glioblastoma multiforme, but, interestingly, the NCI/DTP assays showed temozolomide to be completely inactive (LC_50_ > 100 μM) against the SF-539 cell line. Indeed, the NCI/DTP tests showed temozolomide to be inactive against all 6 CNS cell lines. Thus, **Pt**_**2**_**ad** is significantly more cytotoxic toward the SF-539 cell line than either cisplatin or temozolomide.

**Pt**_**2**_**ad** is also more cytostatic than either cisplatin or temozolomide against the CNS SF-539 cancer cells, as seen by comparing the growth inhibition values in Table 3. For the total growth inhibition (TGI), the concentration of **Pt**_**2**_**ad** need be only 2.76 μM, compared to a required concentration of 7.67 μM for cisplatin. But, for 50% growth inhibition (GI_50_) of a population of SF-539 cells, a cisplatin solution with a concentration of only 0.600 μM is needed, compared to a 1.47 μM solution of **Pt**_**2**_**ad**. Interestingly, for temozolomide, GI_50_ > 100 μM and TGI > 100 μM.

In addition to demonstrating cytotoxicity toward cancer cells *in vitro*, a good chemotherapy drug for treating glioblastoma multiforme must be able to penetrate the blood-brain barrier. The free web tool Swiss ADME^48^ predicts that **Pt**_**2**_**ad** should be able to penetrate the blood-brain barrier, and that the gastrointestinal tract would highly absorb **Pt**_**2**_**ad**. The Swiss ADME calculation also points out the low water-solubility and the high molar mass of **Pt**_**2**_**ad** as limitations on the usefulness of this compound as a chemotherapy drug.

Pt_2_ad was also particularly cytotoxic and cytostatic toward renal cancer cell line RXF-393,^49^ with LC_50_ = 5.99 μM and TGI = 2.63 μM. For this same cell line, LC_50_ = 24.2 μM and TGI = 4.58 μM for cisplatin. Conversely, cisplatin was more cytotoxic and cytostatic than **Pt**_**2**_**ad** against the renal cancer cell line A498. Though notably less cytotoxic toward the ovarian and breast cancer cell lines, **Pt**_**2**_**ad** was nonetheless more cytotoxic than cisplatin toward both the ovarian OVCAR-5 cells and the triple-negative breast cancer cells MDA-MB-231.

## Conclusions

**Pt**_**2**_**ad** joins a growing list of organometallic platinum(IV) derivatives that exhibit anticancer properties.^32, 50, 51, 52, 53, 54, 55^

**Pt**_**2**_**ad** is more cytotoxic than cisplatin against some cell lines of colon cancer (HCT-116, HT29, and SW-620), the glioblastoma multiforme SF-539 cell line, some melanomas (cell lines LOX IMVI, MALME-3M, and SK-MEL-28), the ovarian cancer cell line OVCAR-5, the renal cancer cell line RXF-393, breast cancer cell line BT-549, and triple-negative breast cancer cell line MDA-MB-231. Nevertheless, there were also some cell lines for which cisplatin was more cytotoxic than **Pt**_**2**_**ad**, and a number of cell lines against which **Pt**_**2**_**ad** displayed little to no cytotoxicity at all. When assessing the anticancer properties of a new compound *in vitro*, using as many different cell lines as possible is necessary in order to gauge the chemotherapeutic potential accurately.

## Supporting information

Check CIF Results

Summary of NCI 5-dose results

## Acknowledgments

The authors thank the Department of Chemistry & Biochemistry at the University of Alaska Fairbanks (UAF) for financial support of this work. At UAF, the 600 MHz NMR spectrometer was purchased from funding by the US Army Medical Research and Material Command (05178001), and the 300 MHz NMR spectrometer was purchased from funding by the National Science Foundation (DUE−9850731). Undergraduate research support for A. M. O’B. and support for maintaining the 600 MHz NMR spectrometer at UAF were supplied by an Institutional Development Award (IDeA) from the National Institute of General Medical Sciences of the National Institutes of Health (NIH) under grant number P20GM103395. The content is solely the responsibility of the authors and does not necessarily reflect the official views of the NIH. A. M. O’B. also acknowledges undergraduate research support from the UAF Office of Undergraduate Research and Scholarly Activity (URSA). UAF is an affirmative action/equal employment opportunity employer and education institution: www.alaska.edu/nondiscrimination.) The National Science Foundation’s Major Research Instrumentation is acknowledged for their support (1827313) in the purchase of the Bruker D8 Venture X-ray diffractometer at Whitworth University. The authors thank the National Cancer Institute Developmental Therapeutics Program (NCI/DTP) and acknowledge NCI/DTP (https://dtp.cancer.gov) as the source of the *in vitro* sulforhodamine B assay data for **Pt**_**2**_**ad** (NSC 842139), cisplatin (NSC 119875), and temozolomide (NSC 362856/32).

