## Supplementary material for "Synthesis, Structure and Anticancer Activity of a Dinuclear Organoplatinum(IV) Complex Stabilized by Adenine": Check CIF Results

### checkCIF (basic structural check) running

*Checking for embedded fcf data in CIF ...*
*Found embedded fcf data in CIF. Extracting fcf data from uploaded CIF, please wait* ....

### checkCIF/PLATON (basic structural check)

Structure factors have been supplied for datablock(s) 22wah02a

THIS REPORT IS FOR GUIDANCE ONLY. IF USED AS PART OF A REVIEW PROCEDURE FOR PUBLICATION, IT SHOULD NOT REPLACE THE EXPERTISE OF AN EXPERIENCED CRYSTALLOGRAPHIC REFEREE.

No syntax errors found. CIF dictionary
Please wait while processing .... Interpreting this report

Structure factor report

### Datablock: 22wah02a

```
Bond precision:         C-C = 0.0047 A                  Wavelength=0.71073
Cell:        a=10.1669(9)       b=10.5929(9)       c=13.0266(11)
             alpha=92.574(3)    beta=109.885(3)    gamma=103.182(4)
Temperature: 100 K
```

| | Calculated | Reported |
|---|---|---|
| Volume | 1272.97(19) | 1272.97(19) |
| Space group | P -1 | P -1 |
| Hall group | -P 1 | -P 1 |
| Moiety formula | C11 H23 I2 N5 Pt2, C H Cl3, C2 H6 O | C11 H23 I2 N5 Pt2, C H Cl3, C2 H6 O |
| Sum formula | C14 H30 Cl3 I2 N5 O Pt2 | C14 H30 Cl3 I2 N5 O Pt2 |
| Mr | 1034.74 | 1034.76 |
| Dx,g cm-3 | 2.700 | 2.700 |
| Z | 2 | 2 |
| Mu (mm-1) | 13.732 | 13.732 |

| | | |
|---|---|---|
| F000 | 940.0 | 940.0 |
| F000' | 933.04 | |
| h,k,lmax | 15,16,20 | 15,16,20 |
| Nref | 10216 | 10210 |
| Tmin,Tmax | 0.056,0.404 | 0.389,0.747 |
| Tmin' | 0.008 | |

Correction method= # Reported T Limits: Tmin=0.389 Tmax=0.747 AbsCorr = MULTI-SCAN

Data completeness= 0.999 Theta(max)= 33.778

R(reflections)= 0.0223( 9148) wR2(reflections)= 0.0475( 10210)

S = 1.085 Npar= 263

---

The following ALERTS were generated. Each ALERT has the format
**test-name_ALERT_alert-type_alert-level**.
Click on the hyperlinks for more details of the test.

---

#### Alert level C

PLAT972_ALERT_2_C Check Calcd Resid. Dens. 0.60Ang From Pt1 -2.06 eA-3
**And 5 other PLAT972 Alerts**
More ...
PLAT977_ALERT_2_C Check Negative Difference Density on H1A . -0.35 eA-3
**And 6 other PLAT977 Alerts**
More ...

---

#### Alert level G

PLAT002_ALERT_2_G Number of Distance or Angle Restraints on AtSite 2 Note
PLAT172_ALERT_4_G The CIF-Embedded .res File Contains DFIX Records 1 Report
PLAT232_ALERT_2_G Hirshfeld Test Diff (M-X) Pt1 --I2 . 13.4 s.u.
PLAT860_ALERT_3_G Number of Least-Squares Restraints ............ 1 Note
PLAT883_ALERT_1_G No Info/Value for _atom_sites_solution_primary . Please Do !
PLAT910_ALERT_3_G Missing # of FCF Reflection(s) Below Theta(Min). 1 Note
PLAT912_ALERT_4_G Missing # of FCF Reflections Above STh/L= 0.600 5 Note
PLAT933_ALERT_2_G Number of HKL-OMIT Records in Embedded .res File 1 Note
PLAT978_ALERT_2_G Number C-C Bonds with Positive Residual Density. 2 Info

---

0 **ALERT level A** = Most likely a serious problem - resolve or explain
0 **ALERT level B** = A potentially serious problem, consider carefully
13 **ALERT level C** = Check. Ensure it is not caused by an omission or oversight
9 **ALERT level G** = General information/check it is not something unexpected

1 ALERT type 1 CIF construction/syntax error, inconsistent or missing data
17 ALERT type 2 Indicator that the structure model may be wrong or deficient
2 ALERT type 3 Indicator that the structure quality may be low

2 ALERT type 4 Improvement, methodology, query or suggestion
0 ALERT type 5 Informative message, check

---

It is advisable to attempt to resolve as many as possible of the alerts in all categories. Often the minor alerts point to easily fixed oversights, errors and omissions in your CIF or refinement strategy, so attention to these fine details can be worthwhile. In order to resolve some of the more serious problems it may be necessary to carry out additional measurements or structure refinements. However, the purpose of your study may justify the reported deviations and the more serious of these should normally be commented upon in the discussion or experimental section of a paper or in the "special_details" fields of the CIF. checkCIF was carefully designed to identify outliers and unusual parameters, but every test has its limitations and alerts that are not important in a particular case may appear. Conversely, the absence of alerts does not guarantee there are no aspects of the results needing attention. It is up to the individual to critically assess their own results and, if necessary, seek expert advice.

**Publication of your CIF in IUCr journals**

A basic structural check has been run on your CIF. These basic checks will be run on all CIFs submitted for publication in IUCr journals (*Acta Crystallographica*, *Journal of Applied Crystallography*, *Journal of Synchrotron Radiation*); however, if you intend to submit to *Acta Crystallographica Section C* or *E* or *IUCrData*, you should make sure that full publication checks are run on the final version of your CIF prior to submission.

**Publication of your CIF in other journals**

Please refer to the *Notes for Authors* of the relevant journal for any special instructions relating to CIF submission.

---

**PLATON version of 06/07/2023; check.def file version of 30/06/2023**

### **Datablock 22wah02a** - ellipsoid plot

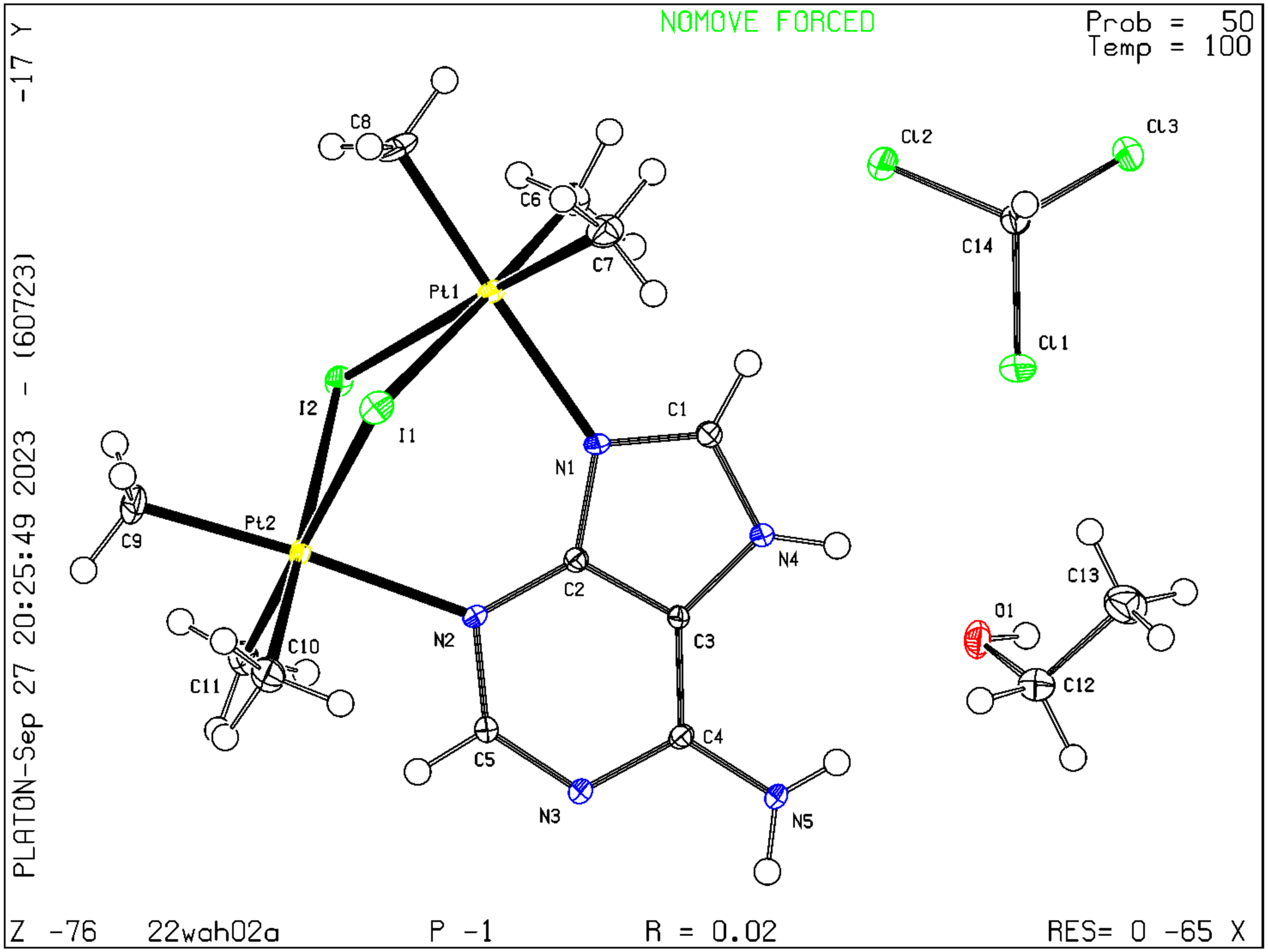


Download CIF editor (publCIF) from the IUCr
Download CIF editor (enCIFer) from the CCDC
Test a new CIF entry
