## Supplementary material for "Synthesis, Structure and Anticancer Activity of a Dinuclear Organoplatinum(IV) Complex Stabilized by Adenine": Summary of NCI 5-dose results

### National Cancer Institute Developmental Therapeutics Program
#### In-Vitro Testing Results

| NSC : D - 842139 / 1 | Experiment ID : 2308NS57 | Test Type : 08 | Units : Molar |
|---|---|---|---|
| Report Date : September 06, 2023 | Test Date : August 14, 2023 | QNS : | MC : |
| COMI : WAHAOB | Stain Reagent : SRB Dual-Pass Related | SSPL : 1CTN | |

| | | | Log10 Concentration | | | | | | | | | | | | |
|---|---|---|---|---|---|---|---|---|---|---|---|---|---|---|---|
| | Time | | Mean Optical Densities | | | | | Percent Growth | | | | | | | |
| Panel/Cell Line | Zero | Ctrl | -8.1 | -7.1 | -6.1 | -5.1 | -4.1 | -8.1 | -7.1 | -6.1 | -5.1 | -4.1 | GI50 | TGI | LC50 |
| **Leukemia** | | | | | | | | | | | | | | | |
| CCRF-CEM | 0.662 | 2.454 | 1.879 | 1.853 | 1.341 | 0.449 | 0.394 | 68 | 66 | 38 | -32 | -40 | 3.11E-7 | 2.86E-6 | > 8.25E-5 |
| HL-60(TB) | 0.694 | 2.398 | 2.232 | 1.990 | 1.749 | 0.538 | 0.541 | 90 | 76 | 62 | -22 | -22 | 1.14E-6 | 4.47E-6 | > 8.25E-5 |
| K-562 | 0.245 | 2.064 | 1.565 | 1.526 | 1.335 | 0.420 | 0.299 | 73 | 70 | 60 | 10 | 3 | 1.30E-6 | > 8.25E-5 | > 8.25E-5 |
| MOLT-4 | 0.604 | 2.232 | 1.594 | 1.696 | 1.529 | 0.654 | 0.525 | 61 | 67 | 57 | 3 | -13 | 1.10E-6 | 1.28E-5 | > 8.25E-5 |
| RPMI-8226 | 0.825 | 2.478 | 2.313 | 2.363 | 2.260 | 1.233 | 0.616 | 90 | 93 | 87 | 25 | -25 | 3.23E-6 | 2.56E-5 | > 8.25E-5 |
| SR | 0.538 | 2.219 | 1.885 | 1.926 | 1.696 | 0.428 | 0.489 | 80 | 83 | 69 | -20 | -9 | 1.34E-6 | 4.87E-6 | > 8.25E-5 |
| **Non-Small Cell Lung Cancer** | | | | | | | | | | | | | | | |
| A549/ATCC | 0.452 | 2.460 | 2.266 | 2.220 | 2.283 | 2.149 | 2.057 | 90 | 88 | 91 | 85 | 80 | > 8.25E-5 | > 8.25E-5 | > 8.25E-5 |
| EKVX | 0.619 | 1.847 | 1.750 | 1.658 | 1.728 | 1.542 | 1.285 | 92 | 85 | 90 | 75 | 54 | > 8.25E-5 | > 8.25E-5 | > 8.25E-5 |
| HOP-62 | 0.589 | 1.924 | 1.800 | 1.867 | 1.797 | 1.757 | 1.773 | 91 | 96 | 90 | 87 | 89 | > 8.25E-5 | > 8.25E-5 | > 8.25E-5 |
| HOP-92 | 1.075 | 1.684 | 1.597 | 1.588 | 1.615 | 0.608 | 0.570 | 86 | 84 | 89 | -43 | -47 | 1.62E-6 | 3.87E-6 | > 8.25E-5 |
| NCI-H226 | 1.069 | 1.757 | 1.734 | 1.693 | 1.716 | 1.430 | 0.573 | 97 | 91 | 94 | 52 | -46 | 8.73E-6 | 2.80E-5 | > 8.25E-5 |
| NCI-H23 | 0.705 | 2.112 | 2.010 | 1.984 | 2.038 | 1.708 | 1.010 | 93 | 91 | 95 | 71 | 22 | 2.21E-5 | > 8.25E-5 | > 8.25E-5 |
| NCI-H322M | 0.766 | 1.980 | 1.889 | 1.791 | 1.700 | 1.590 | 1.237 | 92 | 84 | 77 | 68 | 39 | 3.40E-5 | > 8.25E-5 | > 8.25E-5 |
| NCI-H460 | 0.372 | 3.043 | 3.105 | 3.065 | 3.044 | 2.785 | 0.450 | 102 | 101 | 100 | 90 | 3 | 2.39E-5 | > 8.25E-5 | > 8.25E-5 |
| NCI-H522 | 0.874 | 2.054 | 1.856 | 1.829 | 1.862 | 0.484 | 0.352 | 83 | 81 | 84 | -45 | -60 | 1.51E-6 | 3.70E-6 | 1.86E-5 |
| **Colon Cancer** | | | | | | | | | | | | | | | |
| COLO 205 | 0.544 | 2.269 | 2.150 | 2.215 | 2.409 | 1.867 | 0.385 | 93 | 97 | 108 | 77 | -29 | 1.47E-5 | 4.37E-5 | > 8.25E-5 |
| HCC-2998 | 0.834 | 2.919 | 2.685 | 2.600 | 2.714 | 2.420 | 0.383 | 89 | 85 | 90 | 76 | -54 | 1.31E-5 | 3.17E-5 | 7.68E-5 |
| HCT-116 | 0.274 | 2.478 | 2.332 | 2.291 | 2.242 | 0.842 | 0.086 | 93 | 92 | 89 | 26 | -69 | 3.43E-6 | 1.55E-5 | 5.22E-5 |
| HCT-15 | 0.240 | 2.028 | 1.830 | 1.795 | 1.661 | 1.075 | 0.764 | 89 | 87 | 79 | 47 | 29 | 6.53E-6 | > 8.25E-5 | > 8.25E-5 |
| HT29 | 0.283 | 1.948 | 1.805 | 1.709 | 1.846 | 0.998 | 0.108 | 91 | 86 | 94 | 43 | -62 | 5.99E-6 | 2.12E-5 | 6.36E-5 |
| KM12 | 0.601 | 2.619 | 2.601 | 2.581 | 2.604 | 2.377 | 1.599 | 99 | 98 | 99 | 88 | 49 | 7.98E-5 | > 8.25E-5 | > 8.25E-5 |
| SW-620 | 0.327 | 2.010 | 2.056 | 2.121 | 2.142 | 0.549 | 0.065 | 103 | 107 | 108 | 13 | -80 | 3.37E-6 | 1.14E-5 | 3.92E-5 |
| **CNS Cancer** | | | | | | | | | | | | | | | |
| SF-268 | 0.607 | 2.002 | 2.110 | 2.028 | 2.082 | 1.766 | 0.638 | 108 | 102 | 106 | 83 | 2 | 2.12E-5 | > 8.25E-5 | > 8.25E-5 |
| SF-295 | 0.613 | 2.374 | 2.188 | 2.028 | 1.994 | 1.774 | 1.465 | 89 | 80 | 78 | 66 | 48 | 6.65E-5 | > 8.25E-5 | > 8.25E-5 |
| SF-539 | 0.756 | 2.200 | 2.107 | 2.057 | 2.138 | 0.102 | 0.234 | 94 | 90 | 96 | -87 | -69 | 1.47E-6 | 2.76E-6 | 5.20E-6 |
| SNB-19 | 0.543 | 1.955 | 1.865 | 1.747 | 1.769 | 1.381 | 1.127 | 94 | 85 | 87 | 59 | 41 | 2.72E-5 | > 8.25E-5 | > 8.25E-5 |
| SNB-75 | 1.564 | 2.438 | 2.435 | 2.321 | 2.338 | 2.206 | 1.824 | 100 | 87 | 89 | 73 | 30 | 2.83E-5 | > 8.25E-5 | > 8.25E-5 |
| U251 | 0.429 | 1.924 | 1.860 | 1.743 | 1.815 | 1.603 | 0.479 | 96 | 88 | 93 | 78 | 3 | 1.98E-5 | > 8.25E-5 | > 8.25E-5 |
| **Melanoma** | | | | | | | | | | | | | | | |
| LOX IMVI | 0.525 | 3.025 | 2.842 | 2.740 | 2.543 | 1.396 | 0.183 | 93 | 89 | 81 | 35 | -65 | 3.85E-6 | 1.84E-5 | 5.82E-5 |
| MALME-3M | 0.572 | 1.450 | 1.453 | 1.562 | 1.274 | 0.571 | 0.150 | 100 | 113 | 80 | 0 | -74 | 1.95E-6 | 8.19E-6 | 3.92E-5 |
| M14 | 0.492 | 2.116 | 1.797 | 1.761 | 1.630 | 0.816 | 0.202 | 80 | 78 | 70 | 20 | -59 | 2.08E-6 | 1.48E-5 | 6.34E-5 |
| MDA-MB-435 | 0.686 | 2.626 | 2.581 | 2.567 | 2.513 | 1.534 | 0.127 | 98 | 97 | 94 | 44 | -82 | 6.19E-6 | 1.84E-5 | 4.62E-5 |
| SK-MEL-2 | 1.406 | 2.682 | 2.651 | 2.565 | 2.596 | 2.080 | 1.532 | 98 | 91 | 93 | 53 | 10 | 9.59E-6 | > 8.25E-5 | > 8.25E-5 |
| SK-MEL-28 | 0.625 | 1.663 | 1.617 | 1.584 | 1.464 | 1.060 | 0.079 | 96 | 92 | 81 | 42 | -87 | 5.10E-6 | 1.74E-5 | 4.24E-5 |
| SK-MEL-5 | 0.929 | 3.031 | 2.756 | 2.668 | 2.584 | 1.814 | 1.394 | 87 | 83 | 79 | 42 | 22 | 5.03E-6 | > 8.25E-5 | > 8.25E-5 |
| UACC-257 | 0.962 | 2.370 | 2.271 | 2.201 | 2.188 | 1.783 | 1.254 | 93 | 88 | 87 | 58 | 21 | 1.37E-5 | > 8.25E-5 | > 8.25E-5 |
| UACC-62 | 1.014 | 2.836 | 2.738 | 2.642 | 2.372 | 1.609 | 0.160 | 95 | 89 | 75 | 33 | -84 | 3.18E-6 | 1.57E-5 | 4.20E-5 |
| **Ovarian Cancer** | | | | | | | | | | | | | | | |
| IGROV1 | 0.284 | 1.473 | 1.622 | 1.379 | 1.275 | 0.779 | 0.256 | 113 | 92 | 83 | 42 | -10 | 5.20E-6 | 5.31E-5 | > 8.25E-5 |
| OVCAR-3 | 0.611 | 1.917 | 2.065 | 2.003 | 1.996 | 0.966 | 0.383 | 111 | 107 | 106 | 27 | -37 | 4.24E-6 | 2.17E-5 | > 8.25E-5 |
| OVCAR-4 | 0.750 | 2.088 | 1.978 | 1.954 | 1.891 | 1.417 | 0.778 | 92 | 90 | 85 | 50 | 2 | 8.17E-6 | > 8.25E-5 | > 8.25E-5 |
| OVCAR-5 | 0.671 | 2.004 | 1.815 | 1.703 | 1.708 | 0.893 | 0.073 | 86 | 77 | 78 | 17 | -89 | 2.35E-6 | 1.19E-5 | 3.52E-5 |
| OVCAR-8 | 0.729 | 2.872 | 2.842 | 2.818 | 2.827 | 2.474 | 1.015 | 99 | 97 | 98 | 81 | 13 | 2.39E-5 | > 8.25E-5 | > 8.25E-5 |
| NCI/ADR-RES | 0.454 | 1.705 | 1.661 | 1.631 | 1.642 | 1.168 | 0.407 | 96 | 94 | 95 | 57 | -10 | 1.05E-5 | 5.79E-5 | > 8.25E-5 |
| SK-OV-3 | 0.601 | 1.533 | 1.534 | 1.419 | 1.528 | 1.541 | 1.591 | 100 | 88 | 99 | 101 | 106 | > 8.25E-5 | > 8.25E-5 | > 8.25E-5 |
| **Renal Cancer** | | | | | | | | | | | | | | | |
| 786-0 | 0.667 | 2.416 | 2.218 | 2.221 | 2.369 | 2.089 | 0.427 | 89 | 89 | 97 | 81 | -36 | 1.53E-5 | 4.07E-5 | > 8.25E-5 |
| A498 | 1.515 | 2.366 | 2.427 | 2.439 | 2.545 | 2.506 | 0.609 | 107 | 109 | 121 | 116 | -60 | 1.97E-5 | 3.78E-5 | 7.26E-5 |
| ACHN | 0.493 | 2.037 | 1.962 | 1.978 | 1.925 | 1.239 | 0.265 | 95 | 96 | 93 | 48 | -46 | 7.54E-6 | 2.67E-5 | > 8.25E-5 |
| CAKI-1 | 0.576 | 2.058 | 1.823 | 1.837 | 2.024 | 1.377 | 0.479 | 84 | 85 | 98 | 54 | -17 | 9.40E-6 | 4.76E-5 | > 8.25E-5 |
| RXF 393 | 0.991 | 1.213 | 1.174 | 1.095 | 1.148 | 0.303 | 0.105 | 83 | 47 | 71 | -69 | -89 | . | 2.63E-6 | 5.99E-6 |
| SN12C | 0.556 | 2.199 | 1.971 | 1.846 | 1.717 | 1.590 | 0.728 | 86 | 79 | 71 | 63 | 10 | 1.46E-5 | > 8.25E-5 | > 8.25E-5 |
| TK-10 | 0.786 | 1.654 | 1.530 | 1.476 | 1.684 | 1.568 | 0.668 | 86 | 79 | 104 | 90 | -15 | 1.99E-5 | 5.94E-5 | > 8.25E-5 |
| UO-31 | 0.529 | 1.880 | 1.698 | 1.610 | 1.715 | 0.966 | 0.283 | 87 | 80 | 88 | 32 | -47 | 3.96E-6 | 2.12E-5 | > 8.25E-5 |
| **Prostate Cancer** | | | | | | | | | | | | | | | |
| PC-3 | 0.587 | 2.044 | 1.931 | 1.821 | 1.755 | 1.162 | 0.432 | 92 | 85 | 80 | 39 | -26 | 4.55E-6 | 3.27E-5 | > 8.25E-5 |
| DU-145 | 0.373 | 1.670 | 1.699 | 1.688 | 1.553 | 1.284 | 1.030 | 102 | 101 | 91 | 70 | 51 | > 8.25E-5 | > 8.25E-5 | > 8.25E-5 |
| **Breast Cancer** | | | | | | | | | | | | | | | |
| MCF7 | 0.438 | 2.137 | 2.279 | 2.210 | 2.082 | 1.056 | 0.702 | 108 | 104 | 97 | 36 | 16 | 4.91E-6 | > 8.25E-5 | > 8.25E-5 |
| MDA-MB-231/ATCC | 0.513 | 1.148 | 0.995 | 0.981 | 0.958 | 0.524 | 0.215 | 76 | 74 | 70 | 2 | -58 | 1.62E-6 | 8.79E-6 | 6.02E-5 |
| HS 578T | 1.511 | 2.489 | 2.244 | 2.238 | 2.298 | 1.581 | 1.275 | 75 | 74 | 80 | 7 | -16 | 2.14E-6 | 1.69E-5 | > 8.25E-5 |
| BT-549 | 1.045 | 2.122 | 2.067 | 2.072 | 2.027 | 1.328 | 0.404 | 95 | 95 | 91 | 26 | -61 | 3.56E-6 | 1.65E-5 | 6.12E-5 |
| T-47D | 0.709 | 1.814 | 1.764 | 1.754 | 1.685 | 1.509 | 1.534 | 95 | 95 | 88 | 72 | 75 | > 8.25E-5 | > 8.25E-5 | > 8.25E-5 |
| MDA-MB-468 | 0.908 | 1.621 | 1.530 | 1.514 | 1.488 | 0.544 | 0.299 | 87 | 85 | 81 | -40 | -67 | 1.49E-6 | 3.85E-6 | 1.92E-5 |
